# Consistent, linear phenological shifts across a century of observations in South Korea

**DOI:** 10.1101/2022.09.29.510037

**Authors:** William D. Pearse, Michael Stemkovski, Benjamin R. Lee, Richard B. Primack, Sangdon Lee

## Abstract

The Korea Meteorological Agency (KMA) has monitored flowering dates over the past 100 years for seven economically important woody plant species. This unique dataset is perfect for understanding whether historical patterns of phenological plasticity are breaking down in the face of recent and rapid climate change. Here we show that a scientist armed only 50 years into this study would have been able to predict the phenological shifts of the last 50 years with a high degree of accuracy. This is despite record-breaking warm temperatures and unprecedented early flowering, suggesting consistency in phenological shifts over time.

## Main text

Shifts in the timing of spring flowering events are recognized as one of the clearest and most sensitive ecological indicators of climate change^1^□. Numerous efforts are under way to track yearly variation in flowering phenology^2,3^□, but most ongoing studies began in the last few decades and so lack data on phenology before the onset of anthropogenic climate change. Such studies typically calculate phenological sensitivity as the linear relationship between flowering time and temperature, but there is emerging uncertainty as to whether plants may be reaching the limit of their ability to track temperature^4,5^□. If they are reaching this limit, we would expect to see either non-linearities (e.g., asymptotes) in phenological sensitivity^6^□, or increased variability (or a lack of predictability) in comparison with what were once tight, strong relationships^7^□. Indeed, some have suggested that drivers such as winter chilling and photoperiod will become more prominent in cueing flowering as spring temperatures become less limiting^8^□. To gain an understanding of whether climate change is resulting in different patterns of phenological response, we need to compare recent shifts to historical baseline data from before the onset of rapid anthropogenic climate change.

The Korean Meteorological Agency (KMA) has been recording data on spring flowering time for seven woody plant species for 100 years as of 2022, making it, to our knowledge, the longest continuously running phenological monitoring effort in Asia^9^ (Fig. 1)□. This centennial study offers a unique opportunity to compare phenological trends before and after the onset of rapid climate change. The extremity of recent phenology is stark: during this tenth decade of study, the record for the highest average annual temperature was broken six times and, in 2021 alone, species at nine of the 72 sites exhibited the earliest flowering phenology ever recorded. Overall, species flowered an average 4.1 days earlier for every 1°C of warming, and as much as 5.5 days earlier in the case of apricots (Figure 2). Spring temperature explained the highest amount of variation in flowering, at least twice as much as any other factor, with significant contributions from latitude and whether the site was coastal or inland. But do these data signal that spring temperature sensitivity has changed over time in response to accelerated warming?

**Figure 1.**
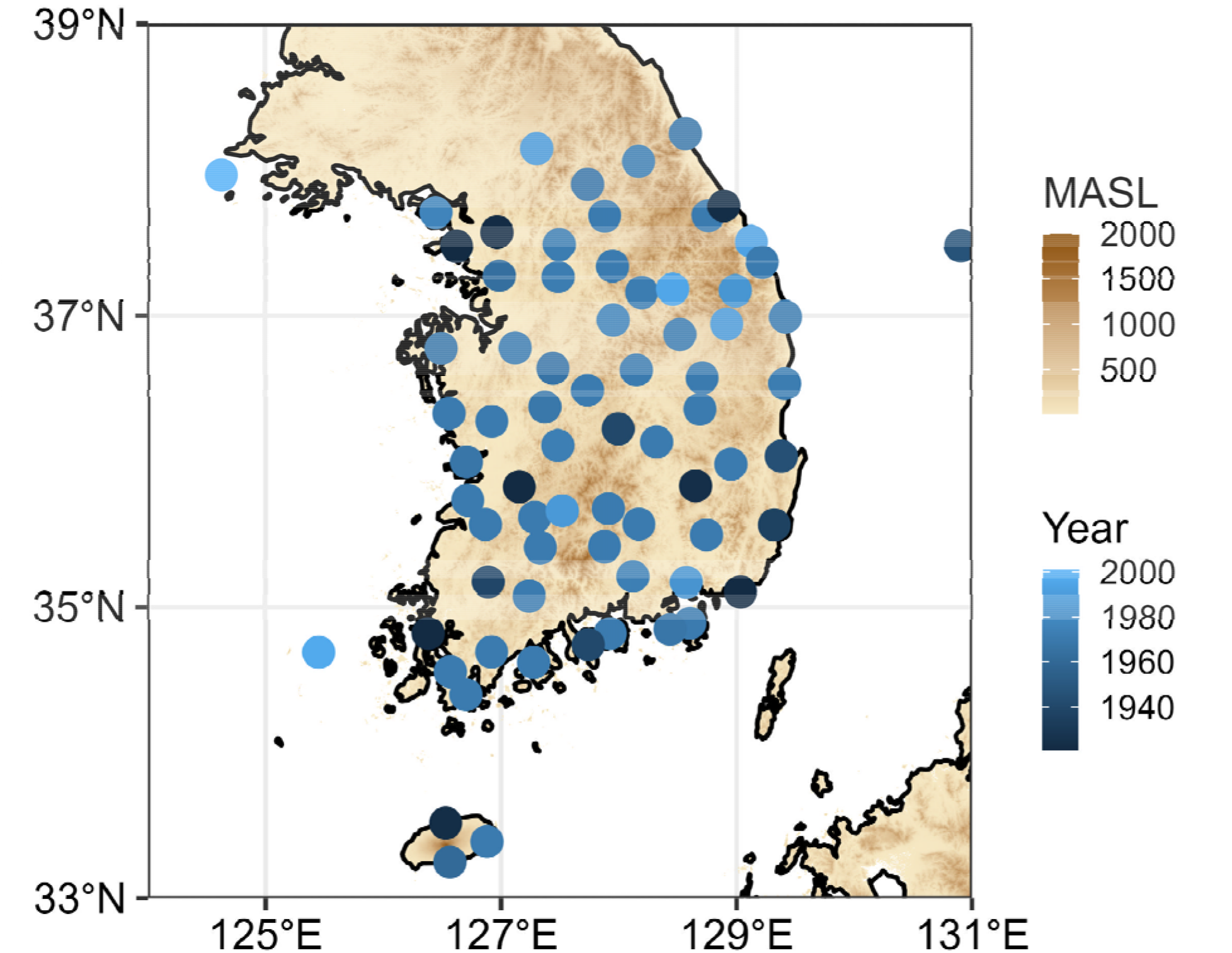
KMA weather station locations used in this study, with point colour indicating the year of first observation. Background colours indicate elevation (meters above sea level).

**Figure 2.**
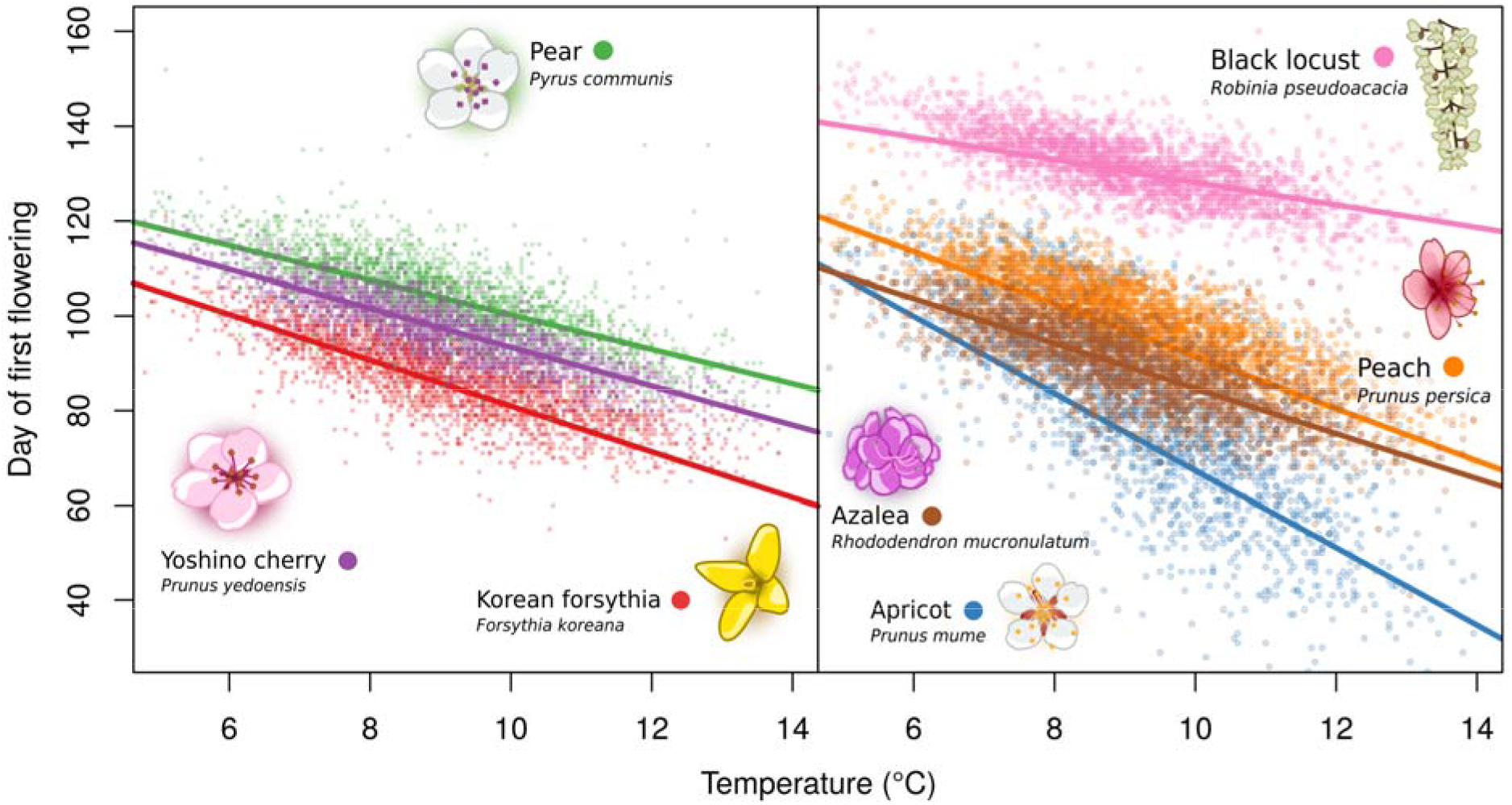
Date of first-flower is consistently dominated by temperature across species. Points represent observations from the second half of the dataset, and the lines represent models fit to the first half of the dataset. All seven species’ date of first flower is plotted against temperature (mean of February-March). Full model summary statistics are given in the supplementary materials, including direct comparisons of the predictive power of temperature vs. other factors (Supplementary Figure S1). Notably, all species’ responses are strongly, linearly correlated with temperature, opening earlier in warmer years.

We evaluated these data with two criteria that could indicate shifting sensitivity to spring temperature: non-linearity in trends and increased inter-annual variability around the mean trend. Regarding linearity, we found that while models were best fit by quadratic and cubic phenology-to-temperature functions for most species, their gains in predictive ability were small, suggesting a high degree of fidelity with linear functions. Averaged across all species, quadratic and cubic models explained <2% more variation than linear models. At most, the error associated with the sensitivity prediction was reduced by only 0.03 days (for forsythia). We found no evidence that an exponential decay response curve was more appropriate than a simple linear fit for any species. Thus, we find little evidence to suggest that the capacity for flowering phenology to respond to spring temperature is saturating in this system (all models givens in Supplementary Materials S3).

We next quantified the degree to which inter-annual phenological variability changed over time. If climate change were disrupting the ability of species to track environmental variability through phenological plasticity, we might expect to see increases in inter-annual variability. This could occur if flowering were to be decoupled from spring temperature through sensitivity saturation, with other factors becoming the dominant drivers^10^. We thus explicitly modeled changes in phenological variance over time as a function of spring temperatures to test for changes in predictability. All but one species (cherry) tended to become, if anything, slightly less variable in their inter-annual phenology over time. Of these, four (forsythia, apricot, black locust, and rhododendron) showed strong evidence (Bayesian p > 95% for estimated slope of σ^2^) that this reduction in variance over time was statistically significant, though we note that these variance reductions are very small when compared to the magnitudes of mean shifts (the greatest shift is in plums, which also have the greatest variance overall; see Supplementary Materials S5). Further, five of the seven species (excepting cherry and pear) showed extremely modest (albeit statistically significant) evidence of reductions in variance in warmer years (model results given in Supplementary Materials S6), suggesting that the reductions in variance over time are not simply a product of shifting temperature variance. These findings indicate that inter-annual phenological variation has not become more variable.

Combined, these two lines of evidence suggest that phenological sensitivity of woody plants in Asia has not significantly changed over the past century. Indeed, it suggests that had scientists in 1971 used the first 50 years of data to predict phenology in 2021, their predictions would have been highly accurate. To quantify this, we modeled the degree to which predictability has changed since the onset of climate change by comparing the predictive accuracy of models trained with the first fifty years and the most recent fifty years of data. A simple linear model containing only one term – average spring temperature – when fit to data from 1921 to 1971, predicts the observed phenology from 1972 to 2021 with an average error of 7.3 days across all species (lowest RMSE is 4.4 days for cherries, highest 13.1 days for apricots). On the other hand, the same model, when fit using the most recent fifty years, had only slightly higher accuracy for all species, with an average error of 6.6 days: a 6.4% improvement. Thus, scientists in 1971 could have made very good predictions of shifts in phenology that would have been accurate for the next 50 years.

These results provide three critical insights for modern climate change research. The first is confirmatory but sometimes overlooked: historic records form an important baseline against which contemporary change must be measured to avoid the problem of shifting baselines^7^. Second, while climate is shifting in unprecedented ways that are challenging to model and forecast, the biological responses to these changes may still be linear and predictable based on existing data. Third, we stress than even these linear phenological responses may still have complex, non-linear, and multifaceted ecological consequences. All species adjust their phenology in response to temperature variation, but their sensitivities differ from one another (note the variation in slopes in Figure 2). For every 1°C of warming, black locust flowering advances by 3.3 days, but apricot advances by 5.5 days. Even subtle, linear differences in species sensitivities have the potential to fundamentally reorder phenological relationships and vulnerability to frost and drought, and profoundly affect ecological interactions and function. This may occur long before species reach hard physiological limits on flowering plasticity. Still, our results suggest that currently available data hold the key to predicting the future consequences of shifts in flowering phenology.

## Supporting information

S5 - variance-year.tsv

S6 - variance-temp.tsv

S3 - response-curves.tsv

S2 - basic-models.tsv

S4 - half-century-predictions.tsv

S1 - korea-pheno-code.zip

## Methods

We analyzed changes in predictability of flowering time in seven species from a 100-year dataset collected by the Korea Meteorological Administration. We describe the data processing and collection briefly due to its long record period and because it is given in more detail elsewhere, and describe the analyses in full detail. All reproducible analysis code are released in the supplementary materials; the data were not collected by us and are housed with the Korea Meteorological Agency (KMA) where they are available on request.

### Data Collection

We used data gathered by the Weather Service of the Republic of Korea on the first-flowering date of *Forsythia koreana* (Korean forsythia), *Prunus mume* (Japanese apricot), *Pyrus communis* (pear), *Prunus × yedoensis* (Yoshino cherry), *Prunus persica* (peach), *Robinia pseudoacacia* (black locust) and, *Rhododendron mucronulatum* (Korean rhododendron) at 74 weather stations distributed across the country, with some weather stations starting in 1921 (data available from Korea Meteorological Administration, http://www.web.kma.go.kr/edu/unv/agri-cultural/seasonob/11733741389.html; Figure 1)^9□^. Monitoring at other weather stations started later, and not all stations have operated continuously. Temperatures have risen across the country over the study period, with greater temperature increases in areas with larger human populations^11□^. These data have been extensively used to investigate how species are changing their phenology over time and in response to annual variation in temperature ^12,13,14^L.

Observations were made on plants growing in phenological gardens on the grounds of each weather station. All plants used in this study were obtained from stock maintained and distributed by the Weather Service of the Republic of Korea to maintain genetic uniformity. When plants died or were not healthy, they were replaced by new plants. To minimize the effects of tree aging, the trees in the phenological garden have been replaced at regular intervals of between 15 and 25 years throughout the study. Surface air temperature data were available as early as from October 1907 (city of Seoul), and records at some stations were missing during the Korean War of 1950-1953. The phenological observations were made by weather station employees according to precise written instructions that have remained constant for the entire study. First flowering was recorded when at least three flowers were open on a plant.

### Statistical analysis

Our analysis was split into four main components: (1) quantifying the drivers of phenological change, (2) assessing changes in forecast skill in recent decades (3) testing for non-linearity in phenological responses, and (4) estimating the predictability of phenology over time and in warmer years.

(1) We chose six predictor variables to explain inter-annual variation in first-flowering phenology: spring temperatures (mean over February and March), spring precipitation (mean over February and March), regional population (https://jumin.mois.go.kr/index.jsp#), elevation, latitude, and whether the station was inland or coastal. While temperature sensitivity is often calculated using degree-day models, we follow a convention in another recent study that was similarly interested in changing responses and variance through time. All continuous predictor variables were centered to have means of 0 and standard deviations of 0.5 to allow for the estimation of standardized effect size and important across continues and discrete variables^15^□. The day-of-year (DOY) of first-flowering was modeled using linear mixed effects models for each species separately, with the predictor variables modeled as additive fixed effects, and station as a random effect with variable intercepts to account for site-specific differences and pseudo-replication. We extracted slope coefficients (fixed effects) from these models to compare the relative importance of the phenology predictors to one another and across species.

(2) In quantifying the drivers of phenology as described above, we found support for temperature as the dominant driver of phenology in this system. As a first test of whether phenological responses to temperature have changed fundamentally over the course of the study, we quantified how surprising phenological patterns in the second half of the dataset (after the onset of climate change; 1972-2021) are compared to what was observed in the first half (pre-climate change; 1921-1971). We did this by assessing the improvement in forecast skill^16^□ that was achieved by the second half of the dataset. For each species, we quantified forecast skill improvement as the percent change of the root-mean-square error (RMSE) of a linear model of DOY as a function of temperature trained on the 1921-1971 data and evaluated on the 1972-2021 data and the RMSE of the same model trained and evaluated on the 1972-2021 data. To obtain one metric of forecast skill improvement, we also calculated the mean of the RMSE estimates across all species.

(3) To assess evidence of non-linearities in species’ responses to temperature, we contrasted models where temperature was modeled in various non-linear ways, using the mean squared-error and AIC of the models as decision criteria. To be both conservative and also maximize the comparability of models (since some non-linear modeling approaches struggle to fit models with as many terms as are present in our models above), we first fit linear (not mixed-effects) models to all explanatory variables in part 1 above, excluding temperature. We then modeled the residuals of these models against: (a) temperature, (b) temperature and its quadratic, (c) temperature and its cubic, and (d) temperature as a saturating exponential decay.

(4) To test for changes in phenological dispersion over time and in warmer years, we follow Pearse et al.^6^ in fitting models which explicitly estimated heteroskedasticity by modeling variance as a function of time and annual temperature. Specifically, we fit Bayesian hierarchical models using *brms*^17^ with the equivalent formulation as in part (1), where the average response (μ) of DOY varies additively as a function of the explanatory, and each station has a hierarchically-drawn estimate for its mean. We then modeled, in two separate models, the variance in response (σ^2^) as a function of time and temperature and the same hierarchical intercept structure to account for variance differences by station. We note that this modelling formulation is conservative because temperature always appears in the μ term, and inclusion of the hierarchical term in both the μ and σ^2^ formulas accounts for changes in station coverage. We used a Gaussian identity link function for the σ^2^ coefficients in order to report variance changes in units of days/year and days/°C. Models were run for 4 chains and 2000 iterations each using default non-informative priors, and were checked for convergence.

## Acknowledgements

We are grateful to XXX anonymous reviewers, and the editorial board, for their help improving this manuscript. WDP and the Pearse lab are also funded by NSF Grant ABI-1759965. SDL are supported by MOE-2020002990006, 2021003360002, and NRF-2021R1A2C1011213.

## Supplementary Materials

**S1 – korea-pheno-code.zip**. Compressed archive of all analysis code.

**S2 – basic-models.tsv**. Outputs from models of all species against all explanatory variables, both scaled (to estimate relative importance of variables) and unscaled.

**S3 – response-curves.tsv**. Outputs from linear and non-linear models to assess support for saturation/non-linearity of species’ functional responses to temperature.

**S4 – half-century-predictions.tsv**. Forecast and hindcast predictions from models, along with within-sample model results.

**S5 – variance-year.tsv**. Outputs from Bayesian hierarchical models with variance terms accounting for correlated changes in variation with temperature.

**S6 – variance-temp.tsv**. Outputs from Bayesian hierarchical models with variance terms allowing for correlated changes in variation with temperature.

## Extended Data

**Figure S1.**
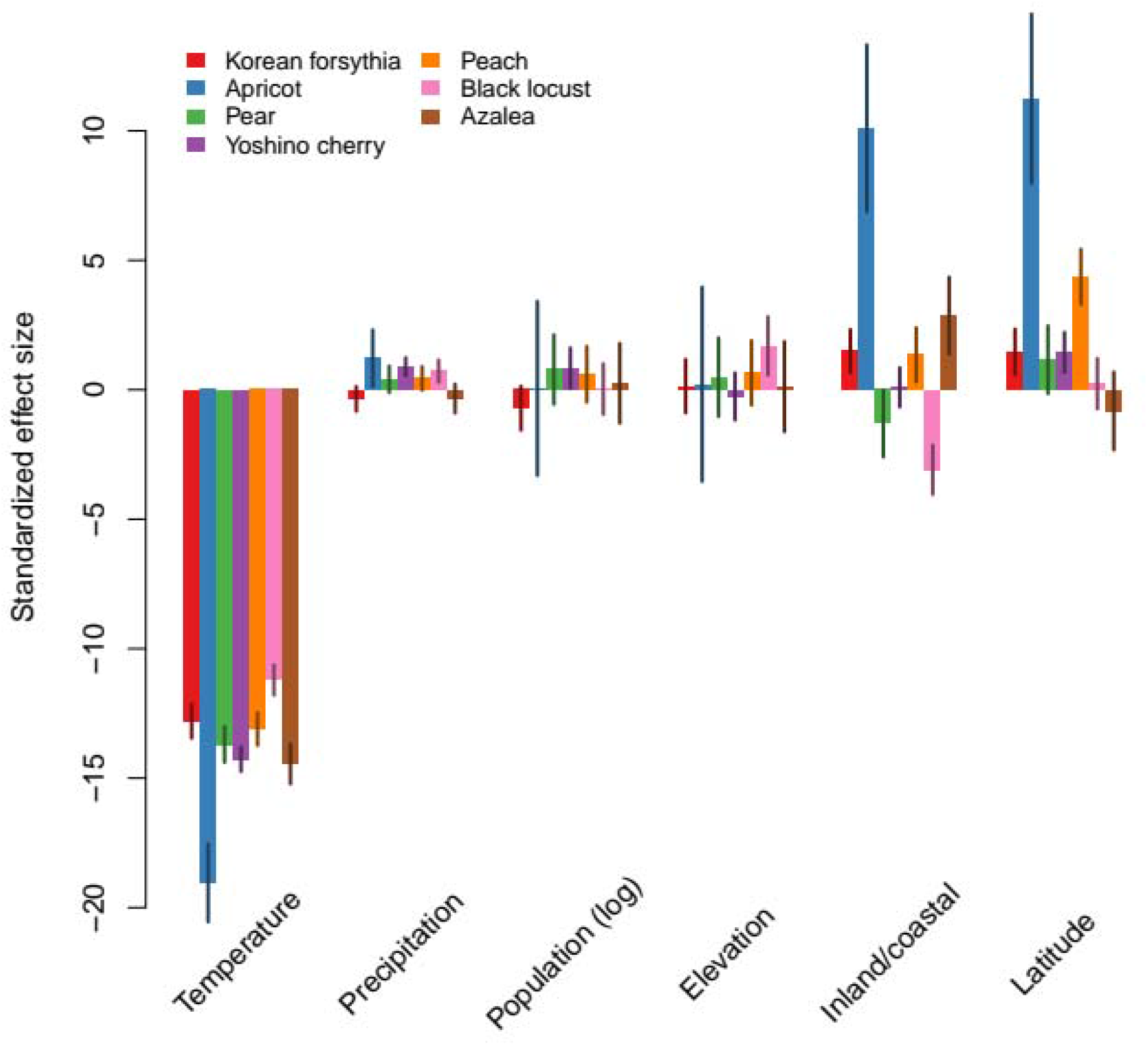
Standardized effect sizes of the drivers of inter-annual variation in flowering phenology. Temperature was the dominant predictor of phenology across all species. Station location also played an important role in predicting phenology, particularly in apricot. Apricots at inland stations tended to flower later than at coastal sites, and most species flower later at higher latitudes.

**Figure S2.**
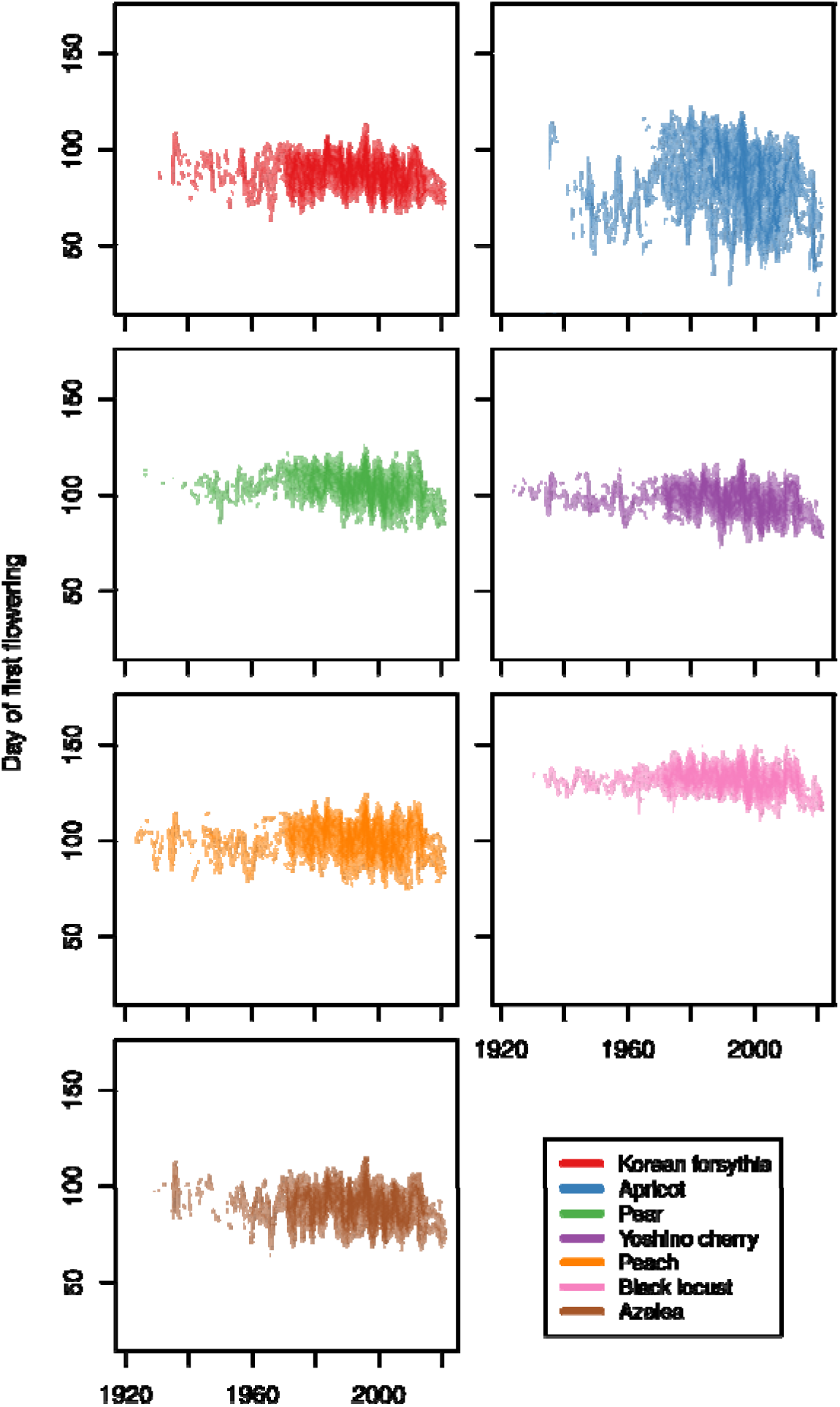
100 years of first-flowering dates in seven species across the Korean Peninsula and islands. Each time-series represents one of 72 sites at which first flowering has been recorded in one of 7 species. At the left-hand side the mean date of first-flower for each species in the first year (1921) is shown, while at the right the mean date for 2021 is shown; transparent horizontal lines map the mean of the 1921 observations per species. We emphasize that not all sites have been monitored for 100 years, and sites joining (and leaving) the network can be seen in the figure.

## Notes

### Competing Interest Statement

The authors have declared no competing interest.

