## Supplementary material for "Consistent, linear phenological shifts across a century of observations in South Korea": S1 - korea-pheno-code.zip: build.sh

```
#!/usr/bin/env bash

########################
# OPTIONAL #############
# Clean previous runs ##
#rm figs-tabs/*
#rm clean-data/*
########################

# Install packages
Rscript src/headers.R
# Clean data
Rscript src/clean-data.R
# Run models (and their table outputs)
Rscript src/models.R
# Make figures
Rscript src/figures.R
```
