## Supplementary material for "Consistent, linear phenological shifts across a century of observations in South Korea": S1 - korea-pheno-code.zip: clean-data.R

```
# Headers
source("src/headers.R")

# Load each sheet
.load.data <- function(sheet){
    print(sheet)
    data <- read.xls("raw-data/phenology data(plant).xlsx", sheet)
    names(data)[c(1, 10:21, 23:34)] <- c("year", paste0("temp.",tolower(names(data)[10:21])), paste0("rain.",tolower(names(data)[23:34])))
    names(data) <- gsub(".1", "", names(data), fixed=TRUE)
    for(i in seq_along(names(data)))
        data[,i] <- as.numeric(data[,i])
    data <- data[,c(1:8, 10:21, 23:34)]
    data$location <- sheet
    names(data) <- tolower(names(data))
    #names(data)[8] <- "forsythia" # shouldn't this be rhododendron?
    names(data)[2] <- "forsythia"
    names(data)[8] <- "rhododendron"
    return(data)
}
data <- do.call(rbind,
                lapply(excel_sheets("raw-data/phenology data(plant).xlsx"), .load.data
                       ))

# Add temperature/precipitation data
data$temp.avg <- apply(data[,11:12], 1, mean, na.rm=TRUE)
data$rain.avg <- apply(data[,23:24], 1, mean, na.rm=TRUE)

# Add site meta-data
site <- read.xls("raw-data/Information.xlsx")
names(site) <- c("location", "pop", "inland", "lat", "long", "altitude")
site$inland <- site$inland=="Inland"
site$pop <- as.numeric(gsub(",", "", site$pop, fixed=TRUE))
data <- merge(data, site, by="location")
data$log.pop <- log10(data$pop)

# look for obvious outliers
hist(data[,"forsythia"], 100)
visreg(lm(forsythia ~ temp.avg, data=data)) # 1 clear outlier
grubbs.test(data[,"forsythia"])

hist(data[,"rhododendron"], 100)
visreg(lm(rhododendron ~ temp.avg, data=data))
grubbs.test(data[,"rhododendron"])

hist(data[,"peach"], 100)
visreg(lm(peach ~ temp.avg, data=data)) # 2 clear outliers
grubbs.test(data[,"peach"])

hist(data[,"plum"], 100)
visreg(lm(plum ~ temp.avg, data=data))
grubbs.test(data[,"plum"])

hist(data[,"cherry"], 100)
visreg(lm(cherry ~ temp.avg, data=data))
grubbs.test(data[,"cherry"])

hist(data[,"acacia"], 100)
visreg(lm(acacia ~ temp.avg, data=data))
grubbs.test(data[,"acacia"])

hist(data[,"pear"], 100)
visreg(lm(pear ~ temp.avg, data=data)) # 1 clear outlier
grubbs.test(data[,"pear"])


# prune 4 outliers
data[which(data$forsythia == max(data$forsythia, na.rm=T)), ]
data[which(data$pear == max(data$pear, na.rm=T)), ]
data[which(data$peach %in% tail(sort(data$peach),2)), ]

data[which(data$forsythia == max(data$forsythia, na.rm=T)), "forsythia"] <- NA
data[which(data$pear == max(data$pear, na.rm=T)), "pear"] <- NA
data[which(data$peach %in% tail(sort(data$peach),2)), "peach"] <- NA

# Make scaled (simplified) data
s.data <- data[,c(1:9,34:41)]
for(i in c(2,10:12,14:17))
    s.data[,i] <- as.numeric(scale(s.data[,i], scale=sd(s.data[,i], na.rm=TRUE)*2))

saveRDS(data, "clean-data/data.RDS")
saveRDS(s.data, "clean-data/s.data.RDS")
```
