## Supplementary material for "Consistent, linear phenological shifts across a century of observations in South Korea": S1 - korea-pheno-code.zip: facts.R

```
# Headers
source("src/headers.R")
data <- readRDS("clean-data/data.RDS")
s.data <- readRDS("clean-data/s.data.RDS")

#######################################
# RECORD BREAKERS #####################
#######################################
# Earliest for each species at each location
#  (non-trivial to functionalise because not long-format data)
n.flower <- earliest.flower <- matrix(nrow=length(sites), ncol=length(species), dimnames=list(sites, species))
warmest.year <- setNames(numeric(length(sites)), sites)
for(i in seq_along(sites)){
    t.data <- data[data$location==sites[i],]
    warmest.year[i] <- t.data$year[which.max(t.data$temp.avg)]
    for(j in seq_along(species)){
        n.flower[i,j] <- sum(!is.na(t.data[,species[j]]))
        if(n.flower[i,j]>0)
            earliest.flower[i,j] <- t.data$year[which.min(t.data[,species[j]])]
    }
}

# Earliest flowering in 2021 stats
prod(dim(earliest.flower))
sum(earliest.flower==2021, na.rm=TRUE)
colSums(earliest.flower==2021, na.rm=TRUE)
length(sites)

# Warmest year in 2021 stats
sum(warmest.year==2021)
length(warmest.year)

# Yearly averages tell a different story
sort(tapply(data$temp.avg, data$year, mean))
years <- years[order(names(years))]
rank(years)
rank(names(years))
```
