## Supplementary material for "Consistent, linear phenological shifts across a century of observations in South Korea": S1 - korea-pheno-code.zip: figures.R

```
# Headers
source("src/headers.R")
data <- readRDS("clean-data/data.RDS")
s.data <- readRDS("clean-data/s.data.RDS")

# Summary stats in the text (harder to do than sounds because of sampling effort changes)
with(data, plot(temp.avg ~ year))
locs <- unique(data$location)
sampled.2021 <- 0
earliest.2021 <- setNames(rep(0, length(species)), species)
warmest.2021 <- setNames(rep(0, length(species)), species)
for(i in seq_along(locs)){
    t.data <- data[data$location==locs[i],]
    if(max(t.data$year)==2021){
        sampled.2021 <- sampled.2021+1
        earliest.2021 <- earliest.2021 +
            as.numeric(sapply(species, function(x) t.data$year[which.min(t.data[,x])]) == 2021)
        warmest.2021 <- warmest.2021 + as.numeric(which.max(t.data$temp.avg)==2021)
        with(t.data[which.max(t.data$temp.avg),], points(temp.avg ~ year, pch=20, col="red"))
    }
}
earliest.2021
warmest.2021

# First half regression with second half data
pdf("figs-tabs/century-regression.pdf", width=9, height = 5)
  par(mfrow=c(1,2), mar=c(5,4,2,0))  
  plot(NA, NA, type="n", axes=FALSE, xlab="Temperature (°C)", ylab="Day of first flowering", ylim=c(30,160), xlim=c(5,14))
  axis(1)
  axis(2)
  for(i in c(1,3,4)){
    abline(lm(data[data$year<=1971,species[i]] ~ data$temp.avg[data$year<=1971]), col=cols[i], lwd=3)
    #abline(lm(data[data$year>1971,species[i]] ~ data$temp.avg[data$year>1971]), col=cols[i], lwd=2, lty=2)
    points(data[data$year>1971,species[i]] ~ data$temp.avg[data$year>1971], col=adjustcolor(cols[i],alpha=0.2), cex=.25, pch=19)
  }
  legend("topright", inset=0.05,
         legend=prnt.spp, col=cols,
         lwd=3, cex=0.75)
  box()
  
  par(mar=c(5,0,2,4))  
  plot(NA, NA, type="n", axes=FALSE, xlab="Temperature (°C)", ylab="Day of first flowering", ylim=c(30,160), xlim=c(5,14))
  axis(1)
  #axis(2)
  for(i in c(2,5,6,7)){
    abline(lm(data[data$year<=1971,species[i]] ~ data$temp.avg[data$year<=1971]), col=cols[i], lwd=3)
    #abline(lm(data[data$year>1971,species[i]] ~ data$temp.avg[data$year>1971]), col=cols[i], lwd=2, lty=2)
    points(data[data$year>1971,species[i]] ~ data$temp.avg[data$year>1971], col=adjustcolor(cols[i],alpha=0.2), cex=.4, pch=19)
  }
  box()
  # title(xlab = "Temperature (°C)",
  #       ylab = "",
  #       outer = TRUE, line = 3)
  #legend(5, 55, legend=prnt.spp[1:4], col=cols[1:4], pch=20, bty="n", cex=1)
  #legend(6.25, 50, legend=prnt.spp[5:7], col=cols[5:7], pch=20, bty="n", cex=1)
dev.off()

# Time-series plots
pdf("figs-tabs/century-series.pdf", height=7, width=4)
  par(mfrow=c(4,2),
      oma = c(5,4,0,0) + 0.1,
      mar = c(0,0,1,1) + 0.1)
  for(i in seq_along(species)){
    with(data, plot(NA, NA, type="n", axes=FALSE, ylab="Day of first flowering", xlab="", xlim=c(1921,2021), ylim=c(20,170)))
    if(i %in% c(1,3,5,7)) axis(2)
    if(i %in% c(6,7)) axis(1)
    axis(1, labels=FALSE)
    axis(2, labels=FALSE)
    box()
    for(loc in unique(data$location)){
      lines(data[data$location==loc,species[i]] ~ data$year[data$location==loc], col=alpha(cols[i],.25))
      
    }
  }
  # for(i in seq_along(species)){
  #     abline(h=mean(data[data$year<=1930,species[i]], na.rm=TRUE), col=alpha(cols[i],.25), lty=1, lwd=3)
  #     axis(2, at=mean(data[data$year<=1930,species[i]], na.rm=TRUE), col=cols[i], col.axis=cols[i], las=2, label="", tcl=-1, lwd=3)
  #     #axis(2, at=mean(data[data$year>2011,species[i]], na.rm=TRUE), col=cols[i], col.axis=cols[i], las=2, font=2, label="", tcl=1, lwd=3)
  #     #axis(4, at=mean(data[data$year<=1930,species[i]], na.rm=TRUE), col=cols[i], col.axis=cols[i], las=2, label="", tcl=-1, lwd=3)
  #     axis(4, at=mean(data[data$year>2011,species[i]], na.rm=TRUE), col=cols[i], col.axis=cols[i], las=2, font=2, label="", tcl=-1, lwd=3)
  # }
  plot(NA,axes=F, ylab="",xlab="", xlim=c(0,1), ylim=c(0,1))
  legend("bottom", inset=0.05,
         legend=prnt.spp, col=cols,
         lwd=3, cex=0.85)
  
  title(xlab = "Year",
        ylab = "Day of first flowering",
        outer = TRUE, line = 3)
  # legend(1920, 160, legend=prnt.spp[1], col=cols[1], pch=20, bty="n", cex=.75, horiz=TRUE)
  # legend(1940, 160, legend=prnt.spp[-1], col=cols[-1], pch=20, bty="n", cex=.75, horiz=TRUE)
dev.off()

# effects size figure
pdf("figs-tabs/effect_sizes.pdf")
  bar_plot <- barplot(ses_int[,-1], beside=T, col=cols[species],border=F,
                      ylab="Standardized effect size", xaxt="n",
                      space=rep(c(3,0,0,0,0,0,0),6), ylim=c(min(ses_int[,-1]-(2*ses_se[,-1])),max(ses_int[,-1]+(2*ses_se[,-1]))))
  arrows(x0 = bar_plot, y0 = ses_int[,-1]-(2*ses_se[,-1]), y1 = ses_int[,-1]+(2*ses_se[,-1]),
         angle=90, code=3, length=0, lwd=2,
         col=rep(adjustcolor(cols,red.f=0.5,green.f=0.5,blue.f=0.5),6))
  text(seq(6, 6+(6*8.25), length.out=6), par("usr")[3]-0.3,
           labels=c("Temperature", "Precipitation", "Population (log)", "Elevation", "Inland/coastal", "Latitude"),
           srt=45, pos=1, xpd=T)
  legend("topleft", inset=0.02,
         legend=prnt.spp, fill=cols, border=F,
         cex=0.8, ncol=2, bty="n")
dev.off()
```
