## Supplementary material for "Consistent, linear phenological shifts across a century of observations in South Korea": S1 - korea-pheno-code.zip: headers.R

```
packages <- c("readxl","gdata","brms","scales","lme4","drc","devtools","parallel","visreg","outliers","RColorBrewer")
for(pkg in packages){
    if(!require(package=pkg, character.only=TRUE))
        install.packages(pkgs=pkg, dependencies=TRUE)
    if(!require(package=pkg, character.only=TRUE))
        stop("Cannot install/load ", pkg)
}

if(!require(package="aomisc", character.only=TRUE))
        install_github("onofriandreapg/aomisc")
    if(!require(package="aomisc", character.only=TRUE))
        stop("Cannot install/load aomisc")


species <-  c("forsythia", "plum", "pear", "cherry", "peach", "acacia", "rhododendron")
prnt.spp <- c("Korean forsythia", "Apricot", "Pear", "Yoshino cherry", "Peach", "Black locust", "Azalea")
cols <- c("cherry"="pink","plum"="purple","forsythia"="yellow","rhododendron"="red","peach"="orange","pear"="forestgreen","acacia"="blue")
cols <- brewer.pal(7,"Set1") #more distinguishable colors
cols[6] <- brewer.pal(8,"Set1")[8] # subbing out yellow for more readable pink
names(cols) <- species
sites <- c("Andong", "Baengnyeongdo", "Boeun", "Bonghwa", "Boryeong", "Buan", "Busan", "Buyeo", "Changwon", "Cheonan", "Cheongju", "Cheorwon", "Chuncheon", "Chungju", "Chupungryeong", "Daegu", "Daegwallyeong", "Daejeon", "Donghae", "Ganghwa", "Gangneung", "Geochang", "Geoje", "Geumsan", "Goheung", "Gumi", "Gunsan", "Gwangju", "Haenam", "Hapcheon", "Heuksando", "Hongcheon", "Icheon", "Imsil", "Incheon", "Inje", "Jangheung", "Jangsu", "Jecheon", "Jeju", "Jeongeup", "Jeonju", "Jinju", "Juam", "Miryang", "Mokpo", "Munkyeong", "Namhae", "Namwon", "Pohang", "Samcheok", "Sancheong", "Seogwipo", "Seosan", "Seoul", "Sokcho", "Sungsan", "Suwon", "Taebaek", "Tongyeong", "Uiseong", "Uljin", "Ulleung", "Ulsan", "Wando", "Wonju", "Yangpyeong", "Yeongcheon", "Yeongdeok", "Yeongju", "Yeongwol", "Yeosu")
```
