## Supplementary material for "Consistent, linear phenological shifts across a century of observations in South Korea": S1 - korea-pheno-code.zip: models.R

```
# Headers
source("src/headers.R")
data <- readRDS("clean-data/data.RDS")
s.data <- readRDS("clean-data/s.data.RDS")

tsv.wrapper <- function(tables, names, file){
    write.table(matrix(c("",names[1]), nrow=2), file, col.names=FALSE, row.names=FALSE)
    write.table(tables[[1]], file, append=TRUE, col.names=NA, row.names=TRUE)
    for(i in seq(2, to=length(tables))){
        write.table(matrix(c("",names[i]), nrow=2), file, col.names=FALSE, row.names=FALSE, append=TRUE)
        write.table(tables[[i]], file, append=TRUE, col.names=NA, row.names=TRUE)        
    }
}

#######################################
# BASIC MODELS ########################
#######################################
raw_int <- ses_int <- ses_slope_int <- raw_se <- ses_se <- ses_slope_se <- matrix(NA, nrow=length(species), ncol=7, dimnames=list(species,c("int","temp","rain","log.pop","altitude","inland", "lat")))
for(i in seq_along(species)){
    # random intercepts by site only
    raw_model <- eval(substitute(lmer(XXX ~ temp.avg + rain.avg + log.pop + altitude + inland + lat + (1|location), data=data), list(XXX=as.name(species[i]))))
    ses_model <- eval(substitute(lmer(XXX ~ temp.avg + rain.avg + log.pop + altitude + inland + lat + (1|location), data=s.data), list(XXX=as.name(species[i]))))
    # random intrecepts by site and random slopes over years
    ses_slope_model <- eval(substitute(lmer(XXX ~ temp.avg + rain.avg + log.pop + altitude + inland + lat + (year|location), data=s.data), list(XXX=as.name(species[i]))))
    
    # coefficients
    raw_int[i,] <- fixef(raw_model)
    ses_int[i,] <- fixef(ses_model)
    ses_slope_int[i,] <- fixef(ses_slope_model)
    
    # standard errors
    raw_se[i,] <- summary(raw_model)$coefficients[,2]
    ses_se[i,] <- summary(ses_model)$coefficients[,2]
    ses_slope_se[i,] <- summary(ses_slope_model)$coefficients[,2]
    
    # visualizing
    par(mfrow=c(2,3))
    eval(substitute(visreg(lmer(XXX ~ temp.avg + rain.avg + log.pop + altitude + inland + lat + (1|location), data=data), main=species[i]), list(XXX=as.name(species[i]))))
    #plot(NA,xlim=c(0,1),ylim=c(0,1), main=species[i])
}

mean(raw_int[,2])

tsv.wrapper(list(raw_int, ses_int, ses_slope_int),
            c("RAW COEF - random intercepts","SES COEF - random intercepts","SES COEF - random intercepts & slopes"),
            "figs-tabs/basic-models.tsv")

tsv.wrapper(list(raw_se, ses_se, ses_slope_se),
            c("RAW COEF - random intercepts","SES COEF - random intercepts","SES COEF - random intercepts & slopes"),
            "figs-tabs/basic-models_se.tsv")

#######################################
# FIRST VS. SECOND 50 YEARS ###########
#######################################
pre.post <- function(sp, data, s.data, pre=NA, train_years=NA, test_years=NA, model_id="temp"){
    
  if(FALSE){
    sp <- "forsythia"
    train_years <- 1921:1971
    test_years <- 1972:2021
    pre <- NA
    model_id <- "full"
  }
  
  if(!(model_id == "temp" | model_id == "full")) stop("Invalid model ID. Only 'temp' and 'full' are allowed")
  
  if(!any(is.na(pre))){
      t.data <- s.data[data$year %in% pre,] # training set
      p.data <- s.data[!data$year %in% pre,] # test set
    } else{
      t.data <- s.data[data$year %in% train_years,] # training set
      p.data <- s.data[data$year %in% test_years,] # test set
    }
  
    t.data$species <- t.data[,sp]; p.data$species <- p.data[,sp]
    
    # model fit to training set
    if(model_id == "temp") model <- lm(species ~ temp.avg, data=t.data)
    if(model_id == "full") model <- lm(species ~  temp.avg + rain.avg + log.pop + altitude + inland + lat, data=t.data) 
    # model evaluated on test set
    p.data$p.species <- predict(model, p.data) 
    # "species" is observed, and "p.species" is predicted
    return(c(
        #model.r2=summary(model)$r.squared, # training set R2
        #pred.r2=cor(p.data$species, p.data$p.species, use="complete.obs")^2 # test set R2
        model.rmse = sqrt(mean(residuals(model)^2)),
        pred.rmse = sqrt(mean((p.data$species - p.data$p.species)^2, na.rm=T))
    ))
}

train1_test2 <- sapply(species, pre.post, data=data, s.data=data,
                       train_years = 1921:1971, test_years = 1972:2021) # first half is training, second half test
train2_test2 <- sapply(species, pre.post, data=data, s.data=data,
                       train_years = 1972:2021, test_years = 1972:2021) # second half training and test
train12_test2 <- sapply(species, pre.post, data=data, s.data=data,
                         train_years = 1921:2021, test_years = 1972:2021) # full dataset training and test
train12_test12 <- sapply(species, pre.post, data=data, s.data=data,
                         train_years = 1921:2021, test_years = 1921:2021) # full dataset training and test

tsv.wrapper(list(train1_test2, train2_test2, train12_test2, train12_test12),
            c("train_first_test_second","train_second_test_second","train_full_test_second","train_full_test_full"),
            "figs-tabs/half-century-predictions-rmse-temp-only.tsv")

mean(train1_test2[2,])
mean(train2_test2[2,])
mean(train12_test2[2,])
mean(train12_test12[2,])

paste("Phenology in the second half of the dataset is explained by temperature",
      round((mean(train1_test2[2,]) - mean(train2_test2[2,]) / mean(train1_test2[2,])), 2),
      "% better by the second half than the first half")

paste("Phenology in the second half of the dataset is explained by temperature",
      round((mean(train1_test2[2,]) - mean(train12_test2[2,]) / mean(train1_test2[2,])), 2),
      "% better by the second half than the first half")

# forecast accuracy improvement with full model
train1_test2_full <- sapply(species, pre.post, data=data, s.data=data, model_id="full",
                       train_years = 1921:1971, test_years = 1972:2021) # first half is training, second half test
train2_test2_full <- sapply(species, pre.post, data=data, s.data=data, model_id="full",
                       train_years = 1972:2021, test_years = 1972:2021) # second half training and test
train12_test2_full <- sapply(species, pre.post, data=data, s.data=data, model_id="full",
                        train_years = 1921:2021, test_years = 1972:2021) # full dataset training and test
train12_test12_full <- sapply(species, pre.post, data=data, s.data=data, model_id="full",
                         train_years = 1921:2021, test_years = 1921:2021) # full dataset training and test
mean(train1_test2_full[2,])
mean(train2_test2_full[2,])
mean(train12_test2_full[2,])
mean(train12_test12_full[2,])
paste("Phenology in the second half of the dataset is explained by temperature",
      round((mean(train1_test2_full[2,]) - mean(train2_test2_full[2,]) / mean(train1_test2_full[2,])), 2),
      "% better by the second half than the first half")
paste("Phenology in the second half of the dataset is explained by all variables",
      round((mean(train1_test2_full[2,]) - mean(train12_test2_full[2,]) / mean(train1_test2_full[2,])), 2),
      "% better by the second half than the full dataset")
tsv.wrapper(list(train1_test2_full, train2_test2_full, train12_test2_full, train12_test12_full),
            c("train_first_test_second","train_second_test_second","train_full_test_second","train_full_test_full"),
            "figs-tabs/half-century-predictions-rmse-full-model.tsv")


#######################################
# LINEAR VS. SATURATING RESPONSE ######
#######################################
poly.fits <- function(sp, d, plot=FALSE){
    .r2 <- function(m) summary(m)$r.squared
    d$species <- d[,sp]
    l <- lm(species ~ temp.avg + rain.avg + year + log.pop + altitude + inland + lat + location, data=d)
    q <- update(l, . ~ . + I(temp.avg^2))
    c <- update(q, . ~ . + I(temp.avg^3))
    
    if(plot){
      # visualizing
      par(mfrow=c(1,3))
      visreg(l, "temp.avg", main=sp)
      visreg(q, "temp.avg", main=sp)
      visreg(c, "temp.avg", main=sp)
    }

    
    return(c(linear.r=.r2(l), quad.r2=.r2(q), cubic.r2=.r2(c),
             linear.aic = AIC(l), quad.aic = AIC(q), cubic.aic = AIC(c)))
}
poly <- sapply(species, poly.fits, d=s.data)

nonlinear.fits <- function(sp, d, plot=FALSE){
    d <- na.omit(d[,c(sp,"temp.avg","rain.avg","year","log.pop","altitude","inland","lat","location")])
    d$species <- d[,sp]
    null <- lm(species ~ rain.avg + year + log.pop + altitude + inland + lat + location, data=d)
    d$resid <- residuals(null)
    linear <-  summary(lm(resid ~ temp.avg, data=d))$sigma
    quad <- summary(lm(resid ~ temp.avg + I(temp.avg^2), data=d))$sigma
    cube <- summary(lm(resid ~ temp.avg + I(temp.avg^2) + I(temp.avg^3), data=d))$sigma
    exp.decay <- summary(drm(resid ~ temp.avg, fct = DRC.expoDecay(), data=d))$rseMat[1]
    
    l <- lm(resid ~ temp.avg, data=d)
    q <- lm(resid ~ temp.avg + I(temp.avg^2), data=d)
    c <- lm(resid ~ temp.avg + I(temp.avg^2) + I(temp.avg^3), data=d)
    e <- drm(resid ~ temp.avg, fct = DRC.expoDecay(), data=d)
    
    if(plot){
      #visualizing
      par(mfrow=c(2,2))
      visreg(l, "temp.avg", main=sp)
      visreg(q, "temp.avg", main=sp)
      visreg(c, "temp.avg", main=sp)
      visreg(e, "temp.avg", main=sp)
    }

    return(c(null=summary(null)$sigma, linear=linear, quad=quad, cube=cube, exp.decay=exp.decay,
             null_aic = AIC(null), linear_aic = AIC(l), quad_aic = AIC(q), cube_aic = AIC(c), null_exp = AIC(e)))
}
nonlinear <- sapply(species, nonlinear.fits, data) # un-scaled data so that sigma is in day units

# identifying which model is selected for each species
poly_best_r2 <- apply(poly[1:3,],2,function(x) rownames(poly)[which(x == max(x))])
poly_best_aic <- apply(poly[4:6,],2,function(x) rownames(poly[4:6,])[which(x == min(x))])
poly <- rbind(poly, gsub("\\..*","",poly_best_r2))
poly <- rbind(poly, gsub("\\..*","",poly_best_aic))
rownames(poly)[7:8] <- c("best_model_r2", "best_model_aic")

nonlin_best_sigma <- apply(nonlinear[1:5,],2,function(x) rownames(nonlinear)[which(x == min(x))])
nonlin_best_aic <- apply(nonlinear[6:10,],2,function(x) rownames(nonlinear[6:10,])[which(x == min(x))])
nonlinear <- rbind(nonlinear, nonlin_best_sigma)
nonlinear <- rbind(nonlinear, gsub("\\_.*","",nonlin_best_aic))
rownames(nonlinear)[11:12] <- c("best_model_sigma", "best_model_aic")

mean(as.numeric(poly["linear.r",]))
mean(as.numeric(poly["quad.r2",]))
mean(as.numeric(poly["cubic.r2",]))

mean(as.numeric(nonlinear["null",]))
mean(as.numeric(nonlinear["linear",]))
mean(as.numeric(nonlinear["quad",]))
mean(as.numeric(nonlinear["cube",]))
mean(as.numeric(nonlinear["exp.decay",]))

tsv.wrapper(list(poly,nonlinear),
            c("polynomial","stepwise-nonlinear"),
            "figs-tabs/response-curves.tsv")

#######################################
# VARIANCE SHIFTS #####################
#######################################
var.change <- function(x){
    d <- s.data
    d$species <- d[,x]
    print(x)
    
    # original
    # yr  <- brm(bf(species ~ temp.avg + rain.avg + log.pop + altitude + inland + lat + (year|location), sigma ~ year), data = d)
    # tmp <- brm(bf(species ~ temp.avg + rain.avg + log.pop + altitude + inland + lat + (year|location), sigma ~ temp.avg), data = d)

    yr  <- brm(bf(species ~ temp.avg + rain.avg + log.pop + altitude + inland + lat + (1|location), sigma ~ year + (1|location), family=brmsfamily("gaussian", link_sigma = "identity")), data = d, cores=4)
    tmp <- brm(bf(species ~ temp.avg + rain.avg + log.pop + altitude + inland + lat + (1|location), sigma ~ temp.avg + (1|location), family=brmsfamily("gaussian", link_sigma = "identity")), data = d, cores=4)

    return(list(year=fixef(yr), temp=fixef(tmp)))
}

variance <- sapply(species, var.change)
#saveRDS(variance, "figs-tabs/variance.RDS") # backup
variance <- readRDS("figs-tabs/variance_rand_loc.RDS")

tsv.wrapper(apply(variance, 2, function(x) x[1]), species, "figs-tabs/variance-year.tsv")
tsv.wrapper(apply(variance, 2, function(x) x[2]), species, "figs-tabs/variance-temp.tsv")

# visualizing variance shifts
viz.var.change <- function(x){
  d <- s.data
  d$species <- d[,x]
  model_full  <- lmer(species ~ temp.avg + year + rain.avg + log.pop + altitude + inland + lat + (1|location), data = d)
  model_min  <- lmer(species ~ temp.avg + year + (1|location), data = d)
  
  par(mfrow=c(2,2))
  visreg(model_full, "year", main=x)
  visreg(model_full, "temp.avg")
  visreg(model_min, "year")
  visreg(model_min, "temp.avg")
  
  return()
}

sapply(species, viz.var.change)
```
