## Supplementary figures and images for "Consistent, linear phenological shifts across a century of observations in South Korea"

### flowers.png

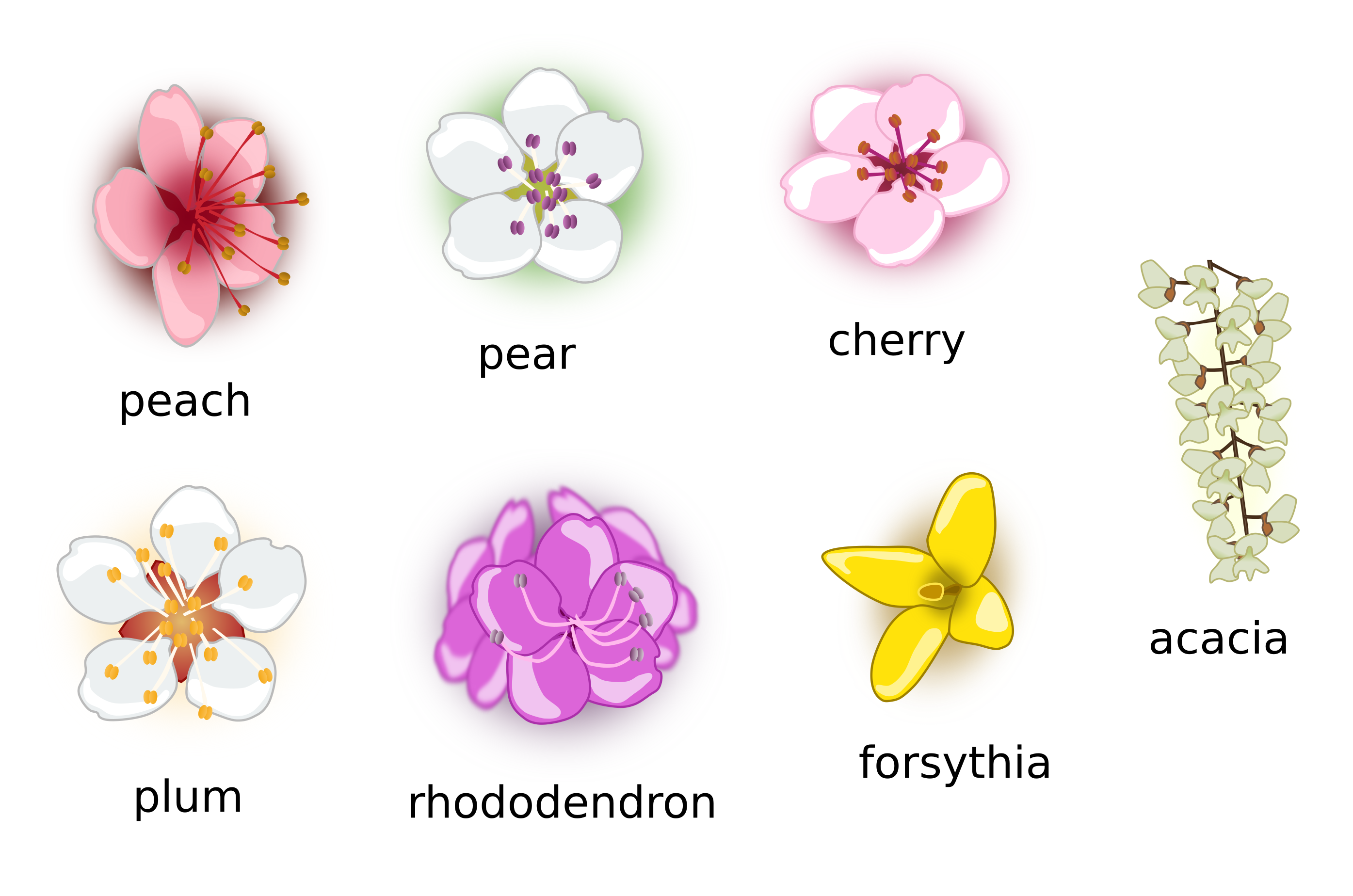
